# Vigilance and behavioral state-dependent modulation of cortical neuronal activity throughout the sleep/wake cycle

**DOI:** 10.1101/2021.06.22.449480

**Authors:** Aurélie Brécier, Mélodie Borel, Nadia Urbain, Luc J. Gentet

**Affiliations:** Integrated Physiology of Brain Arousal Systems, Lyon Neuroscience Research Center, INSERM U1028-CNRS UMR5292, Université Claude-Bernard Lyon 1, 69372 Lyon, France

## Abstract

GABAergic inhibitory neurons, through their molecular, anatomic and physiological diversity, provide a substrate for the modulation of ongoing cortical circuit activity throughout the sleep-wake cycle. Here, we investigated neuronal activity dynamics of parvalbumin (PV), vasoactive intestinal polypeptide (VIP) and somatostatin (SST) neurons in naturally-sleeping head-restrained mice at the level of layer 2/3 of the primary somatosensory barrel cortex of mice. Through calcium-imaging and targeted single-unit loose-patch or whole-cell recordings, we found that PV action potential (AP) firing activity was largest during both NREM (non-rapid eye movement) and REM sleep stages, that VIP neurons were activated during REM sleep and that the overall activity of SST neurons remained stable throughout the sleep/wake cycle. Analysis of neuronal activity dynamics uncovered rapid decreases in PV cell firing at wake onset followed by a progressive recovery during wake. Simultaneous local field potential (LFP) recordings further revealed that, except for SST neurons, a large proportion of neurons were modulated by ongoing delta and theta waves. During NREM sleep spindles, PV and SST activity increased and decreased, respectively. Finally, we uncovered the presence of whisking behavior in mice during REM sleep and show that the activity of VIP and SST is differentially modulated during awake and sleeping whisking bouts, which may provide a neuronal substrate for internal brain representations occurring during sleep.

## INTRODUCTION

The sleep-wake cycle is characterized by specific patterns of cortical activity: high amplitude, low frequency EEG activity in non-rapid eye movement (NREM) sleep and high frequency and low amplitude activity during both wake and rapid-eye movement (REM) sleep. Several hypotheses have emerged as to the function of sleep, with recent evidence pointing to its role in memory consolidation (Born et al., 2006; Girardeau et al., 2009; Vaz et al., 2020) or synaptic homeostasis, a process by which overall neuronal excitability is reduced during sleep (Tononi and Cirelli, 2009; Cirelli, 2017). Delta (1-4 Hz) and spindles (10-17 Hz) oscillations are well known hallmarks of NREM sleep supporting memory formation and consolidation (Steriade and Timofeev, 2003; Maingret et al., 2016; Ulrich, 2016; Latchoumane et al., 2017), whereas theta activity (5-9 Hz) is a marker of exploratory behavior and of REM sleep in rodents (Buzsaki, 2002; Montgomery et al., 2008; Girardeau et al., 2009). It is clear that the internal brain state, and the way sensory cortex processes information are dramatically altered during these vastly different vigilance states, but the underlying circuit mechanisms remain poorly understood.

Cortical networks are composed of both excitatory projection neurons and local inhibitory interneurons, which represent a sparse yet diverse family of GABAergic neurons with different morphologies and electrophysiological properties (Markram et al., 2004; Tremblay et al., 2016), and associated with distinct functional roles in local cortical circuits (Atallah et al., 2012; Gentet et al., 2012; Lee et al., 2013). Despite this diversity, three main subtypes of interneurons have been identified in the past decade: parvalbumin (PV)-containing neurons, vasoactive intestinal peptide (VIP)- and somatostatin (SST)-expressing cells. While there is accumulating evidence that pyramidal cells firing rate evolves over the time course of the sleep/wake cycle (Hobson and McCarley, 1971; Steriade et al., 2001; Watson et al., 2016; Miyawaki et al., 2019), changes in the activity of these three subtypes of interneurons is still poorly understood.

Recent studies have nevertheless revealed brain state-dependent modulation of neocortical interneuron firing activity in different brain regions, notably in the primary somatosensory barrel cortex (S1) during active whisking (Gentet et al., 2010, 2012; Lee et al., 2013; Muñoz et al., 2017). In the motor cortex, additional evidence points toward a vigilance state-dependent of modulation of PV neurons throughout the sleep/wake cycle (Niethard et al., 2016), while SST neurons in the frontal cortex have been shown to be strongly phase-locked to local slow waves sleep oscillations (Funk et al., 2017). In addition, multi-unit recordings, revealed changes in the firing rates of fast-spiking, putative PV cells during NREM and REM sleep in rodents (Vyazovskiy et al., 2009; Vijayan et al., 2010; Watson et al., 2016; Miyawaki et al., 2019) and humans (Peyrache et al., 2011). Finally, a recent study highlighted that acetylcholine and serotonin could both directly modify firing patterns of S1 VIP neurons (Prönneke et al., 2020).

Here, we characterized the activity of the three main subtypes of GABAergic neurons involved in regulating the local excitatory-inhibitory balance in layer 2/3 of S1 in naturally-sleeping, head-restrained mice, using transgenic mouse lines where a red fluorophore could be expressed in genetically-identified subpopulations of GABAergic neurons (Taniguchi et al., 2011). This allowed us to record local cortical neuronal activity across the sleep/wake cycle, either by using calcium imaging, or by targeting PV, VIP and SST neurons under two-photon microscopy visual guidance (Margrie et al., 2003). Both techniques uncovered cell type- and vigilance state-specific differences in neuronal activity, and their results converge to show in both cases increased PV cell activity during NREM and REM sleep, increased VIP cell activity specifically during REM sleep, but an overall stable activity of SST neurons throughout the sleep/wake cycle. Further state-, whisking- and oscillation-dependent shifts in firing patterns and AP firing frequencies revealed the important regulatory role played by GABAergic cells throughout the various stages of the sleep/wake cycle.

## MATERIALS AND METHODS

### Transgenic Mice

Heterozygous male offsprings of GAD2-ires-Cre driver mice (010802, Jackson) crossed with Ai32 (RCLChR2 (H134R)/EYFP) reporter mice (012569, Jackson) were chosen for our experiments.

All animal experiments were conducted after approval by the local ethical committee of the University of Lyon and the Ministry of Science (Protocol number: Apafis #4613) in accordance with the European directive 86/609/EEC on the protection of animals used for experimental and other scientific purposes. All animals used in this study were maintained on a C57Bl6 genetic background and group-housed in the vivarium under normal light cycle conditions.

We used heterozygous 6 – 12 weeks old male and female offsprings of either, PV-ires-Cre mice (008069, Jackson), VIP-ires-Cre (010908, Jackson) or SST-ires-Cre (013044, Jackson) driver mice crossed with Ai9 loxP-tdTomato reporter mice (007909, Jackson). 15 and 60 mice were used in total for the calcium-imaging and patch-clamp studies, respectively. Experiments were performed during the light period of the 12hr : 12hr light-dark cycle. All surgical procedures were performed under isoflurane anesthesia (4 % induction, 1.5-2 % maintenance). A s.c. injection of carprofen (5 mg / kg) and dexamethasone (0.08 mg) was administered prior to surgery to reduce pain and swelling.

First, animals were implanted with a light-weight head-bar allowing head-restraint under two-photon microscopy. The recording chamber was positioned above S1 barrel cortex (−2 mm posterior to bregma and −3.5 mm lateral to midline). Stainless-steel screws and wires were inserted respectively over the contralateral skull (Frontal electrode: +2 mm anterior to bregma, +2 mm lateral; parietal electrode: at −2 mm posterior to bregma, +2 mm lateral) and into the neck muscles for monitoring of EEG and EMG. The skull positions for the EEG screws were chosen to facilitate NREM sleep detection (Fang et al., 2009). After recovery from surgery, mice were habituated to the head-restraint for 3 weeks (daily sessions of increasing durations) until regular bouts of NREM and REM sleeps were detected.

Three days after implantation for the calcium imaging study and on the first day of recordings for the patch clamp study, mice were anesthetized with ~1.5 % isoflurane and the C2 barrel column of the primary somatosensory cortex (S1) was located on the skull using intrinsic optical imaging (Q-Imaging). In short, all whiskers contralateral to the metal head holder chamber were removed except C2. The skull thickness under the chamber was softly reduced with a dental drill and a drop of ASCF was placed inside the chamber. The C2 whisker was deflected at 8 Hz, with an amplitude of 5° for 5 seconds. Successive trials of stimulation were succeeding by trials with no stimulation. Images were processed and analyzed using V++ Software (DigitalOptics, Auckland, New Zealand). The evoked hemodynamic signal was imaged under red LED (610 nm) by subtracting non-stimulation trials to stimulation trials. Finally, the combination of the picture taken under white light and the one taken under red LED, allowed the targeting of C2 column position inside the chamber (Bouchard et al., 2009).

### Virus Injection

For the calcium imaging study, animals were injected with a GCamP6m (AAV1.Syn.GCaMP6m.WPRE.SV40 – Penn Vector Core) (Chen et al., 2013), under isoflurane anesthesia at 1.5-2%, following the intrinsic optical imaging session. A small craniotomy (diameter 100μm) was drilled −1mm lateral to the identified C2 column, and the pia was removed. An injection at 200-300μm depth was performed with a glass pipette inclined at 45° and filled with the virus (0.2μL / min, 2μL delivered in total).

At the end of the habituation period, a small craniotomy (centered on C2 column (~1 x 2 mm) was performed above the C2 barrel column. Dura mater was removed, and 1.5% agarose was applied onto the cortical surface to stabilize it. For the calcium imaging study, a cover slip of the precise dimension of the craniotomy was glued on the skull (Goldey et al., 2014). For patch-clamp experiments, an opening was left on one side of the craniotomy for pipette insertions. The exposed brain was continually immersed in ASCF. Animals were allowed to recover for at least 2-3h before recording, and recording sessions lasted up to 4h. Animals were placed on the stage of a LaVision 2-photon microscope combined with a pulsed Ti-Sapphire laser (λ= 920 nm; Chameleon; Coherent). Image and data acquisition was obtained using Imspector Pro (LaVision, Germany). Simultaneous LFP and whole-cell/cell-attached recordings were performed using glass micropipettes filled respectively with ACSF (in mM: 135 NaCl, 5 KCl, 5 HEPES, 1 MgCl_2_, 1.8 CaCl_2_, 0.01 Alexa-488 (adjusted to pH 7.3 with NaOH) and internal solution (in mM: 135 potassium gluconate, 4 KCl, 10 HEPES, 10 phosphocreatine, 4 MgATP, 0.3 Na3GTP, 0.01 Alexa-488 (adjusted to pH 7.3 with KOH; osmolarity adjusted to 300 mOsmol).

### Calcium Imaging

Images (256 x 91 pixels, 1 pixel = 1.76 μm^2^) were acquired at a frequency of 10.12 Hz. Every recording lasted 2, 3 or 4 minutes. The recording onset was determined by the experimenter to maximize the chances of obtaining bouts of NREM, REM sleep and wakefulness. One animal could be recorded up to 15 consecutive days. A custom-made Matlab routine (Jorrit Montijn, Universiteit van Amsterdam and Mélodie Borel, Université Claude Bernard de Lyon) was used to quantify the GCaMP6-related fluorescence variations of neurons. In short, small x-y drifts; were corrected with an image registration algorithm. If a substantial z-drift was detected during a session, that session was rejected and removed from further analysis. Regions of interest (neuronal somata, and neuropil) were determined semi-automatically using our custom-made Matlab software and ΔF/F0 values for all neurons were then calculated as follows: each neuron’s fluorescence (F) was normalized by the baseline fluorescence of that neuron (F0), taken as the mean of the lowest 50% fluorescence for that session (ΔF = (F - F0)/F0) (adapted from Goltstein et al., 2013). Neurons that were also expressing TdTomato were subsequently categorized as PV, VIP or SST cells, while all other neurons were considered to be putative excitatory pyramidal cells.

### Patch-Clamp/LFP Recordings

Glass pipettes (open tip resistances: 3-4 MOhms for LFP pipettes, 4-6 MOhms for loose-patch or whole-cell patch-clamp pipettes) were lowered into the C2 column of S1 barrel cortex under positive pressure, using two micro-manipulators (Scientifica PatchStar, United Kingdom). Once a neuron of interest was identified in the field of view, the LFP pipette was positioned nearby (~ 100 – 200 μm away). Loose-patch recordings were obtained by approaching the neuron under low pressure until the open tip resistance increased to ~100 MOhms. Whole-cell recordings were obtained by perforating the membrane through rapid suction after obtaining a gigaohm seal. Z-stack images were performed to confirm the position of the pipette tip relative to the neuron of interest. Signals were acquired at 20 kHz and low-pass Bessel filtered at 8 kHz with a Multiclamp 700 B amplifier (Molecular Devices, Axon Instruments). Spikes were; automatically detected using a manually-set threshold on the derivative of the signal.

Oscillatory bouts (Delta waves: 1 – 4 Hz; theta oscillations: 5 - 9 Hz; Spindles: 10 – 17 Hz) were detected using scripts adapted from the Freely Moving Animal (FMA) Matlab Toolbox (developed by M. Zugaro, Collège de France). After band-pass filtering the LFP signal in the appropriate frequency range, a low-pass filtered squared envelope of the z-scored signal was computed. Only bouts that crossed the threshold of z = 2, contained a peak of at least z = 5 and lasted at least 250 ms and 110 ms for delta and theta oscillations respectively, were considered. Bouts were further merged if separated by less than 500ms. Finally, only episodes lasting over 1 s for delta and 350 ms for theta, corresponding to a minimum of 3 oscillations per bout, were kept. Spindles were detected by filtering LFP signal between 10 and 17 Hz and the same low-pass filtered than previously described have been computed. All event has to cross a threshold of 2.5 with a minimum peak of 4. Too close events (less than 200ms) were merged. Finally, only events lasting over 500 ms and less than 3.5 s were kept. Delta events preceding spindles were manually detected after filtering the LFP signal between 0.5 to 4 Hz (minimum and maximum durations of 100 ms and 800 ms respectively). Finally, for the analysis of neuronal activity during delta-spindles bouts, the peak of the delta event was used to align all delta-spindles bouts.

On each detected oscillation, a Hilbert transform was applied and the phase of each spike relative to the oscillation was computed. Only cells with more than 30 spikes detected on oscillations was taken into account for this analysis. All circular statistics were performed using the Circular Statistics Toolbox developed on Matlab (Berens et al., 2009). First, a circular test (Rayleigh’s test) was performed to determine if cells were significantly modulated by the oscillation (p<0.01). Then, a Von Mises distribution was fitted on each spike distribution of modulated cells and two parameters were extracted: the preferred phase (μ) and the concentration (κ). The Von Mises μ corresponds to the mean location of the peak of the distribution, while the Von Mises κ estimates how much the distribution is concentrated on μ.

### Sleep scoring and whisker tracking

EEG and EMG signals were recorded at 20 kHz, amplified (AM system Inc, Model 3000 AC/DC, and band-pass filtered (EEG: 0.1-100 Hz, EMG: 300-3000 Hz) for online display during recording sessions. Sleep scoring was performed manually offline with at a temporal precision of 100 ms. Parameters used to help in the identification of the three main vigilance states (Wake, NREM and REM) were EEG and EMG variance, theta/delta ratio and an ongoing FFT. Wake episodes were defined by a low EEG variance and a high EMG variance. NREM sleep bouts were determined based on a lower EMG and higher EEG variance, together with the presence of spindles on the EEG and/or the LFP. REM sleep was scored when a high theta/delta ratio was observed on the EEG, accompanied by a very low EMG variance. Episodes with undefined vigilance states, such as transitional and intermediate states or episodes lasting less than 20 s were removed from the analysis. Whisker movement was tracked with a high speed (320×240 pixels; 100 Hz) camera (MotionProRedlake, USA) under infrared illumination (λ: 850 nm; OPT machine Vision). The angle of whisker deflection was calculated using a custom-made software written in Matlab (courtesy of P-A. Libourel). An episode of free whisking was considered as such if the C2 whisker deflection exceeded 20 degrees and contained at least three protraction-retractions exceeding 10 degrees, lasting at least 200 ms. Twitches were also detected (> 20 degrees but less than three protraction-retractions exceeding 10 degrees detected) and were removed from the analysis of wakefulness without whisking. Under these conditions, free whisking episodes were detected in both wake and REM sleep, but not during NREM sleep.

### Statistics

All classical statistical analysis was performed using routine functions in Matlab. Normality distribution and homoscedasticity was checked on samples using the Kolmogorov-Smirnov and Levene test respectively. In each case, samples were nonparametric and medians ± median absolute deviations are stated in the section Results. A Wilcoxon-sign ranked test and Friedman test were applied for 2-paired groups and multiple group comparison respectively (unless otherwise stated). If compared samples were considered to be independent, a Mann-Whitney test, for two group comparison, or a Kruskal Wallis test, for multiple groups comparison, were carried out. Finally, when multiple groups comparison appeared significantly different, corrected Wilcoxon-sign ranked tests and corrected Mann-Whitney tests were performed two by two for paired and independent data respectively. For patch-clamp experimental data, some neurons were only recorded in a subset of the three main vigilance states (for example, wakefulness and NREM sleep, but not REM sleep). Therefore, samples were analyzed with a linear mixed-effects model (*LMM*) on R studio using LME4 package and Satterthwaite approximation (Bates et al., 2015). For a better fit to data, repeated measurements were included. In this case, the estimate means and standard errors, computed by Kenward-Roger approximation, are stated in the section Results. All statistical significances are represented by asterisks: * for p < 0.05, ** for p < 0.01, *** for p < 0.001.

## RESULTS

### Cell-specific and vigilance state-dependent calcium activity throughout the sleep-wake cycle

We first investigated how the activity of excitatory and inhibitory neurons might evolve throughout the sleep-wake cycle using *in vivo* two-photon calcium imaging. Transgenic mice expressing Cre tdTomato in either PV, VIP or SST cells, were injected with a calcium indicator (AAVsyn-GCaMP6m), allowing us to specifically identify and measure fluorescence changes in both interneurons and putative pyramidal cells in layers 2/3 of S1 barrel cortex (Fig. 1*A, B*). In total, 2338 putative pyramidal (ie: neurons unlabeled with TdTomato), 310 PV, 113 VIP, 184 SST cells were recorded but only 1206 putative pyramidal, 164 PV, 78 VIP and 134 SST were recorded in all three main vigilance states of the sleep-wake cycle and considered for analysis (Wake, NREM sleep and REM sleep, n = 15 mice). Overall, we observed subtype-specific changes in activity throughout the different vigilance states (Fig. 1*C*). Notably, PV activity increased during sleep stages compared to wake (ΔF/F0: Wake: 11.9 ± 8.0 %; NREM: 18.1 ± 9.1 %; REM: 20.2 ± 10.6 %, n = 164; p < 0.001 for Wake vs NREM and Wake vs REM) and the activity of VIP cells was significantly larger in REM sleep compared to NREM (ΔF/F0, Wake: 37.4 ± 22.9 %; NREM: 25.6 ± 14.3 %; REM: 57.1 ± 34.9 %, n = 78; p < 0.001 for NREM vs REM). The average activity of both putative PYR and GABAergic SST neurons remained however unchanged throughout the sleep-wake cycle (Fig. 1*C, D*). Subdividing the population of PYR neurons into sextiles based on their average activity during wake revealed a larger activity variance during REM compared to wake and NREM sleep (σ: Wake = 70.1, NREM = 70.6, REM = 79.7, *Levene’s test* p < 0.001, Fig. 1*D*). This observation prompted us to separate PYR neurons into two groups based on their average activity during REM sleep. High REM activity PYR displayed significantly lower activity in wake and NREM sleep (ΔF/F0, Wake: 52.2 ± 28.3 %; NREM: 47.3 ± 25.1 %; REM: 71.2 ± 27.0 %, n = 603; p < 0.001 for both Wake vs REM and NREM vs REM) whereas low REM activity PYR cells showed increased wake and NREM activity (ΔF/F0, Wake: 25.3 ± 11.6 %; NREM: 25.3 ± 11.1 %; REM: 18.3 ± 7.5%, n = 603; p < 0.001 for both Wake vs REM and NREM vs REM) (Fig. 1*E*). Furthermore, the average wake and NREM activity of high REM activity PYR cells was larger than their low REM activity counterparts (*Mann-Whitney test*, p < 0.001) (Fig. 1*E*).

**Figure 1:**
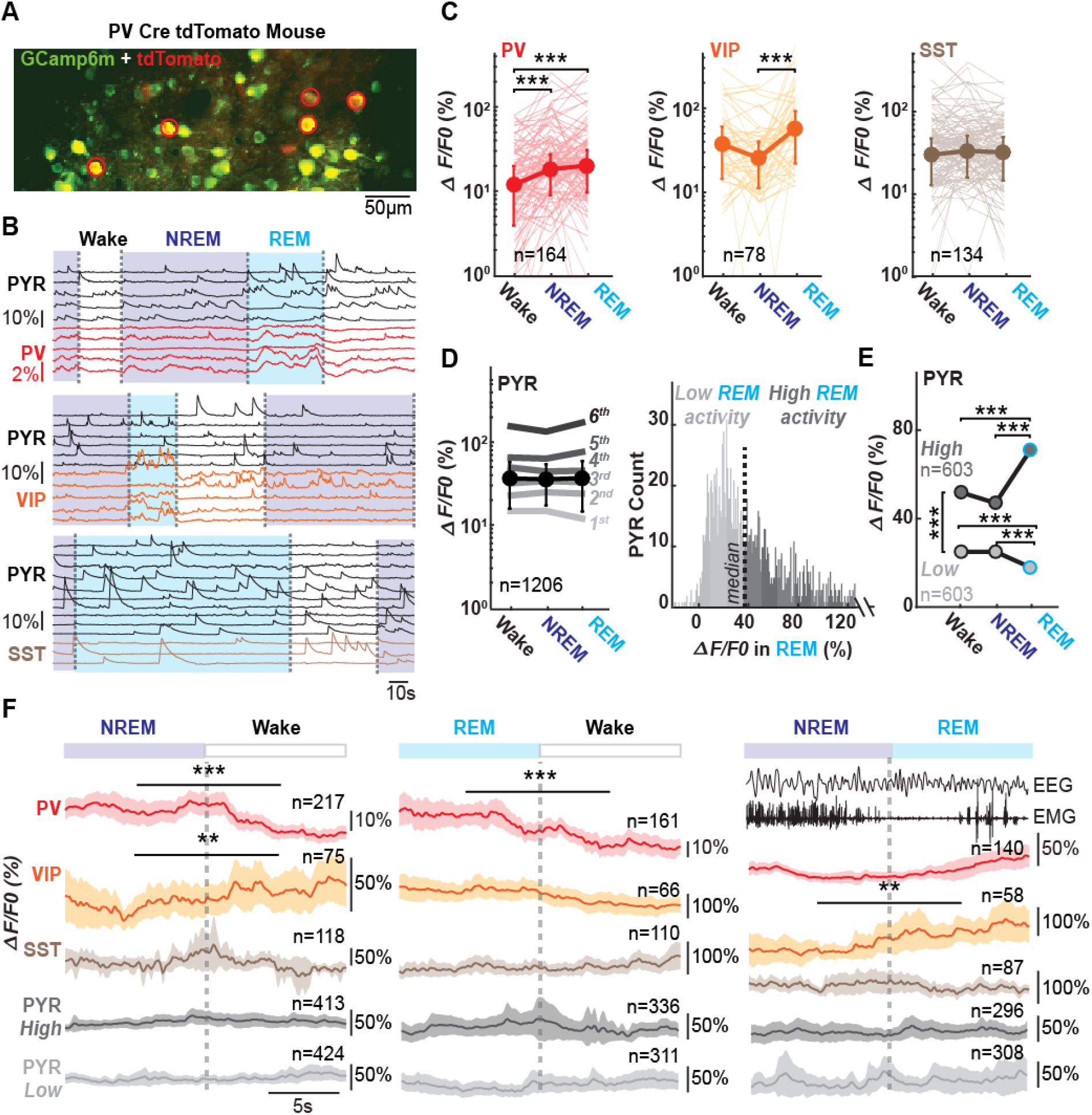
Calcium fluorescence activity changes across the sleep-wake cycle for pyramidal cells and interneurons. **A.** Example of a recording field in the layers 2/3 of barrel cortex in a PV Cre-tdTomato mouse. All neurons in green express GCaMP6m, while PV neurons express in addition tdTomato in red. PV cells expressing both red and green fluorescences are marked by a red circle. **B.** Changes in GCaMP6m fluorescence across time and vigilance states for each detected neurons in one recording field. Example calcium traces in a PV Cre-tdTomato (top, same as in A), a VIP Cre-tdTomato (middle) and in a SST Cre-tdTomato mouse (bottom). **C.** Median changes in fluorescence across vigilance states for each interneuronal subtype. **D.** Median changes in fluorescence of putative PYR across vigilance states (bold line). Additionally, PYR cell population was divided into sextiles according to their activity during wake (left). Histogram distribution of pyramidal cells according to their activity in REM sleep, divided separated into two halves, PYR with low REM activity and PYR with high REM activity (right). **E.** Changes in fluorescence for high REM and low REM activity PYR across vigilance states. **F.** Dynamics of calcium activity at wake onset from NREM (left), REM (middle) sleep, and at REM onset from NREM sleep (right, from −10 s to +10 s around transitions marked by a dotted grey line). Above right, Example raw EEG and EMG traces for a NREM to REM transition. *PV: parvalbumin cells; VIP: vaso-actif intestinal peptide cells; SST: somatostatin cells; PYR: pyramidal cells. Medians and median absolute deviations are represented in C and D and Friedman tests were performed. Medians are represented in E and Friedman tests were performed to compare activity during vigilance state while Kruskall Wallis tests were performed to compare activity of PYR low and high REM activity. Means and confidence intervals are represented in F and Wilcoxon sign rank tests were performed.*

Next, we decided to examine the dynamics of calcium activity around vigilance state transitions. Transitions between vigilance states were manually detected and aligned at t = 0 s (Fig. 1*F*). Awakenings from both NREM and REM sleep stages provide a good temporal resolution through abrupt increases in EMG activity. While the transition between NREM and REM sleep is often considered progressive, we were able, in some instances, to time-stamp such transitions when clear theta waves began to appear on the EEG, concomitant with a rapid reduction in EMG activity and the appearance of heartbeats (Fig. 1*F*). The mean calcium activity between −10 to 0 s from the transition was compared to the activity between 0 to + 10 s. We found that calcium activity of PV and VIP cells decreased and increased respectively at NREM to wake onsets (PV: NREM to Wake transitions: 15.1 ± 11.1 % vs 10.4 ± 8.2 %, n = 217 cells, p < 0.001; VIP: NREM to Wake transitions: 16.0 ± 14.5 % vs 26.6 ± 19.6 %, n = 75 cells, p = 0.004). In addition, PV neurons activity also decreased at wake onset after a REM episode while VIP cell calcium fluorescence increased at REM onset (PV: REM to Wake transitions: 9.3 ± 15.3 % vs −1.3 ± 12.3 %, n = 161 cells, p < 0.001; VIP: NREM to REM transitions: 16.8 ± 22.5 % vs 39.0 ± 40.3 %, n = 58 cells, p = 0.003) (Fig. 1*F*). There was no change in calcium fluorescence in SST neurons and in both groups of PYR cells around any considered state. Taken together, our results suggest that changes in vigilance states occurring throughout the sleep-wake can rapidly alter circuit dynamics through PV and VIP cells of layer 2/3 S1 barrel in naturally-sleeping head-fixed mice.

### Vigilance state-dependent modulation of GABAergic neurons firing activity

While the dynamics of calcium fluorescence across the sleep-wake cycle are a good indicator of overall activity changes, their temporal resolution remains limited. In order to better understand the circuit activity changes induced by different brain vigilance states, we next performed targeted loose- and whole-cell patch-clamp recordings of genetically-identified PV, VIP and SST neurons, as well as putative excitatory pyramidal cells under two-photon microscopy (PV loose-patch n = 32, whole-cell n = 3; VIP loose-patch, n = 31, whole-cell n = 3; SST loose-patch n = 32, whole-cell n = 5 including one silent; PYR loose-patch n = 18, whole-cell n = 4 including one silent; Fig. 2*A, B*), while simultaneously recording the local field potential (LFP) with a second glass pipette positioned in layer 2/3 of the C2 barrel column. In the next section, estimated means ± estimated standard errors of mean are stated and firing activity was compared over the three vigilance states using LMM (see Methods).

**Figure 2:**
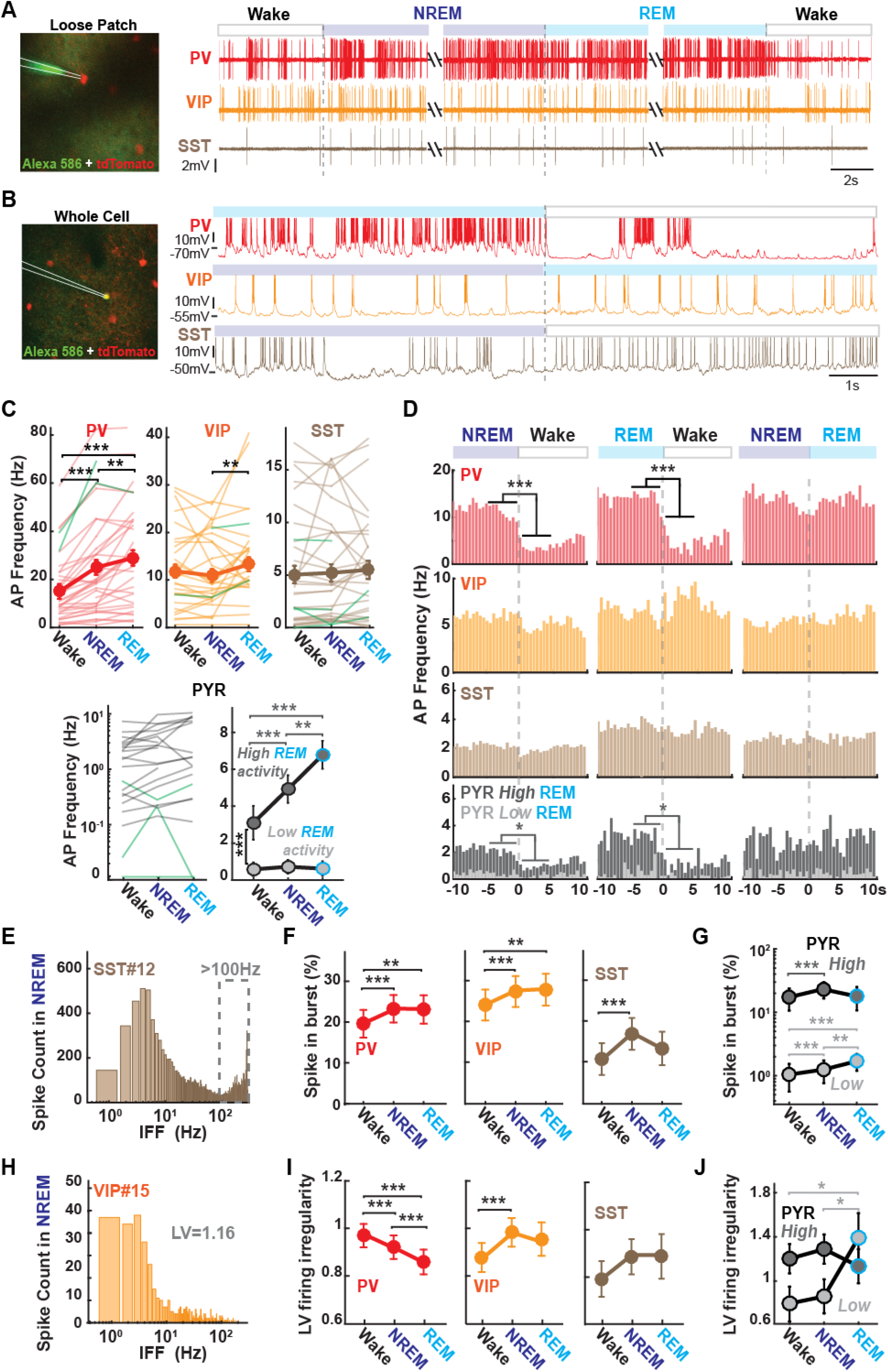
Sleep-wake cycle influences cortical firing rates and pattern. **A.** Loose-patch recordings. Left, z-stack image illustrating the position of a glass pipette (white lines) in loose-patch configuration. Right: example raw traces for a PV, VIP and SST neuron across different vigilance states. **B.** Whole-cell patch-clamp recordings. Left, z-stack image illustrating the position of a glass pipette (white lines) in whole-cell-patch-clamp configuration. Right, example membrane potential traces for a PV, VIP and SST neuron across different vigilance states. APs have been truncated. **C.** Top: Average firing activity for each interneuronal subtype across vigilance states (bold). Individual loose-patched cells ae represented in light and whole-cell patched neurons in green. Bottom left: Firing rates for putative PYR cells across vigilance states in loose-patch (in grey) or whole-cell (green) configurations. Bottom right: Mean AP frequencies for high and low REM activity PYR cells. **D.** Average histograms for different vigilance state transitions (bin size = 500 ms) for each neuronal subtype. **E.** Example instantaneous firing frequency (IFF) distribution for one representative SST cell during NREM sleep. Frequency range > 100 Hz was chosen to define high-frequency bursts. **F.** Percentage of high frequency bursts according to vigilance state for PV, VIP and SST cells. **G.** Same as in F, but for putative PYR neurons with high and low REM activity. **H.** Same as in E for one representative VIP cell during NREM sleep. Local coefficient of variation (LV) measures of the firing irregularity have been estimated for each cell and vigilance states. **I.** LV for each neuronal cell type across vigilance states. **J.** Same as in I, but for PYR cells with high and low REM activity. *AP: action potential. Estimated means and standard errors of mean are represented in bold in C, F, G, I and J and LMM were performed (see Methods). Means are represented in D and Wilcoxon sign rank tests were performed.*

AP firing activity of PV cells increased in NREM compare to wake (p < 0.001) and increased in REM sleep compared to both wake (p < 0.001) and NREM (p = 0.003) (Wake: 15,0 ± 3.0 Hz, n = 34; NREM: 25.0 ± 3.0 Hz, n = 35; REM: 29.0 ± 3.2 Hz, n = 31; *LMM*) (Fig. 2*C*). Firing rates of VIP cells significantly increased during REM sleep compared to NREM (p = 0.002), mirroring our results found under calcium-imaging (Wake: 11,9 Hz ± 1.4, n = 31; NREM: 10.9 Hz ± 1.4, n = 31; REM: 13.3 Hz ± 1.5, n = 28; *LMM*). Finally, the AP firing activity of SST cells remained unchanged overall across vigilance states. For PYR neurons, we split our populations into two groups as previously described, after removing a subset of cells that were not recorded during REM sleep (n = 18 cells). On average, high REM activity PYR were less active in wake compared to NREM and REM (Wake: 3.1 ± 0.6 Hz, n = 9; NREM: 4.9 ± 0.7 Hz, n = 9; REM: 6.8 ± 0.9 Hz, n = 9, p < 0.001 for Wake vs NREM, p < 0.001 for Wake vs REM, p = 0.007 for REM vs NREM; *LMM*) while low REM activity PYR cells did not display any significant changes in overall AP firing activity across vigilance (Fig. 2*C*). The activity of these two subgroups of PYR neurons was also consistently significantly different across all vigilance states (*Mann-Whitney test* p < 0.001 for Wake, NREM and REM) (Fig. 2*C*). Taken together, these findings are consistent with our calcium activity results (Fig. 1), and confirm that PV cells activity increases during NREM and REM sleep compared to wake, that VIP activity specifically increases during REM sleep compared to NREM, and that SST neuronal activity is overall not affected by the different stages of the sleep-wake cycle.

As for our calcium-imaging study, we were able to manually time-stamp clear transition points between vigilance states, both at wake onset, but also between NREM and REM sleep in some instances. Therefore, we investigated the dynamics of AP firing rates of different neuronal cell types around those transition time points (aligned at t = 0 s) (Fig. 2*D*). We quantified and compared firing rates from −5 s to 0 s and from 0 to +5 s. Confirming the results we found under calcium-imaging, a rapid decrease in PV cell firing was observed at wake onset from either NREM or REM sleep (NREM to Wake transitions: 8.5 ± 5.2 Hz vs 1.0 ± 1.0 Hz; n = 32, p < 0.001; REM to Wake transitions: 12.7 ± 9.2 Hz vs 2.0 ± 1.2 Hz; n = 30, p < 0.001; *LMM*) and no changes in the firing rate of SST cells and PYR low REM activity were observed around transitions. However, we did not observe any fast changes in VIP cells firing across states transitions. Finally, firing rates of PYR high REM activity cells decreased significantly as soon as the mice woke up, which was not revealed by calcium imaging data (NREM to Wake transitions: 2.3 ± 0.7 Hz vs 1.5 ± 2.3 Hz; n = 9, p = 0.04; REM to Wake transitions: 2.0 ± 3.0 Hz vs 0.0 ± 4.0 Hz; n = 9, p = 0.016; *LMM*).

While cortical neurons are often classified as bursty on non-bursty on the basis of a clear bimodality in their firing ISIs (Peyrache et al., 2011), we observed that most of our recorded interneurons could fire at both high and low frequency regimes throughout the sleep-wake cycle, as previously observed for other neuronal types, for example in the thalamus (Urbain et al., 2019). In particular, high frequency bursts can strongly modulate synaptic integration and could therefore increase information content (Lisman, 1997). Therefore, we chose to investigate whether GABAergic and putative excitatory neurons firing patterns changed throughout the sleep-wake cycle by focusing on high-frequency bursts firing, defined as any instance of an instantaneous firing frequency (IFF) falling above 100 Hz for VIP, SST and PYR neurons, or rising above 200 Hz for PV cells (Fig. 2*E*). For this analysis, only episode containing more than 30 spikes were taken into account. On average, the proportion of high frequency bursts in PV and VIP neurons increased in both NREM and REM sleep compared to wake (PV, Wake: 19.6 ± 3.4%, n = 34; NREM: 23.2 ± 3.4 %, n = 35; REM: 23.1 ± 3.5 %, n = 31; p < 0.001 between Wake and NREM, p = 0.018 for Wake vs REM; VIP, Wake: 24.2 ± 3.8 %, n = 31; NREM: 27.5 ± 3.8 %, n = 31; REM: 27.9 ± 3.9 %, n = 28; p < 0.001 for Wake vs NREM, p = 0.008 for Wake vs REM; *LMM*). In contrast, SST cells, as well as PYR neurons with high REM activity, displayed a more bursting mode specifically in NREM sleep compared to wake (SST Wake: 10,9 % ± 3.9, n = 29; NREM: 17.1 ± 3.9 %, n= 30; REM: 13.5 ± 4.1 %, n = 26; p < 0.001 for Wake vs NREM; High REM PYR, Wake: 17.3 ± 6.6 %; NREM: 23.2 ± 6.6 %; REM: 17.9 ± 7.2 %, n = 9; p = 0.01 between Wake and NREM; *LMM*) (Fig. 2*F, G*). On the other hand, PYR neurons with low REM activity, showed a significant increase in their bursting behavior in NREM and further more during REM sleep (Low REM PYR, Wake: 1.0 ± 0.5 %; NREM: 1.3 ± 0.5 %; REM: 1.7 ± 0.5 %, n = 9; p = 0.01 between Wake and NREM, p < 0.001 for Wake vs REM; p = 0.002 for NREM vs REM; *LMM*) (Fig. 2*G*). These observed shifts in high-frequency bursts firing may be related to changes in the firing irregularity of neurons. To quantify this parameter, we calculated the local coefficient of variation (LV) for which an LV of 1, more than 1 or less than 1, indicates Poisson-like, irregular and regular firing respectively (Vinck et al., 2016) (Fig. 2*H*). Interestingly, we found that LV firing irregularity was vigilance-state and cell-type specific. PV firing was significantly more irregular in wake, while VIP firing was more irregular in NREM sleep (PV, Wake: 0.97 ± 0.05, n = 34; NREM: 0.92 ± 0.05, n = 35; REM: 0.86 ± 0.052, n = 31, p < 0.001 for both Wake vs NREM and Wake vs REM and NREM vs REM; VIP, Wake: 0.87 ± 0.06, n = 31; NREM: 0.98 ± 0.06, n = 31; REM: 0.95 ± 0.07, n = 28, p < 0.001 for NREM vs Wake; *LMM*) and SST cells displayed the same firing pattern across vigilance state (Fig. 2*I*). In contrast, the firing irregularity of PYR low REM activity increased significantly in REM sleep compared to wake and NREM (Low REM PYR: Wake: 0.8 ± 0.2; NREM: 0.9 ± 0.2; REM: 1.4 ± 0.2, n = 9; p = 0.01 for Wake vs REM; p = 0.03 for NREM vs REM; *LMM*), while firing pattern of high REM PYR neurons remained unchanged across all vigilance states (Fig. 2*J*).

### Theta and delta oscillations modulate timing of interneuronal firing

We observed a variety of oscillations on the layer 2/3 LFP recorded via a glass pipette positioned nearby. LFP cortical oscillations in the theta band (5-9 Hz) have been previously observed during both wakefulness and REM sleep, but not during NREM sleep (Montgomery et al., 2008; Del Vechio Koike et al., 2017). Interestingly, we could also distinguish prominent theta oscillations during REM sleep on our layer 2/3 LFPs, although the LFP electrode is usually positioned in deeper layers (Fig. 3*C*). We therefore decided to investigate phase locking of local interneurons and putative pyramidal cells to layer 2/3 LFP theta oscillations detected in wake and REM sleep (Fig. 3*A - C*). First, we estimated that 58% and 48 % of PV cells were significantly modulated by theta in wake and REM respectively (Rayleigh test, see Methods), 60 % and 43 % of VIP cells, 41 % and 16 % of SST cells, and 47 % and 44 % of PYR neurons (Fig. 3*D*). Fits using von Mises functions yielded estimates of the preferred phase (μ) and the concentration (κ) of spiking (which represents how peaked the spiking distribution is around the preferred phase – See Methods) (Fig. 3*B*). Interestingly, all recorded cells seems to fire at the trough of the theta oscillation, regardless of their neuronal identities and the vigilance state (Fig. 3 *E, F*). On the other hand, the median concentration parameter κ of PV cells during wake remained higher than the one obtained during REM sleep (p = 0.03) while the median κ was similar for the other cells in both wake and REM sleep (PV κ: Wake: 0.88 ± 0.2, n = 19; REM: 0.54 ± 0.15, n = 14, *corrected Mann-Whitney test*) (Fig. 3*E, G*).

**Figure 3:**
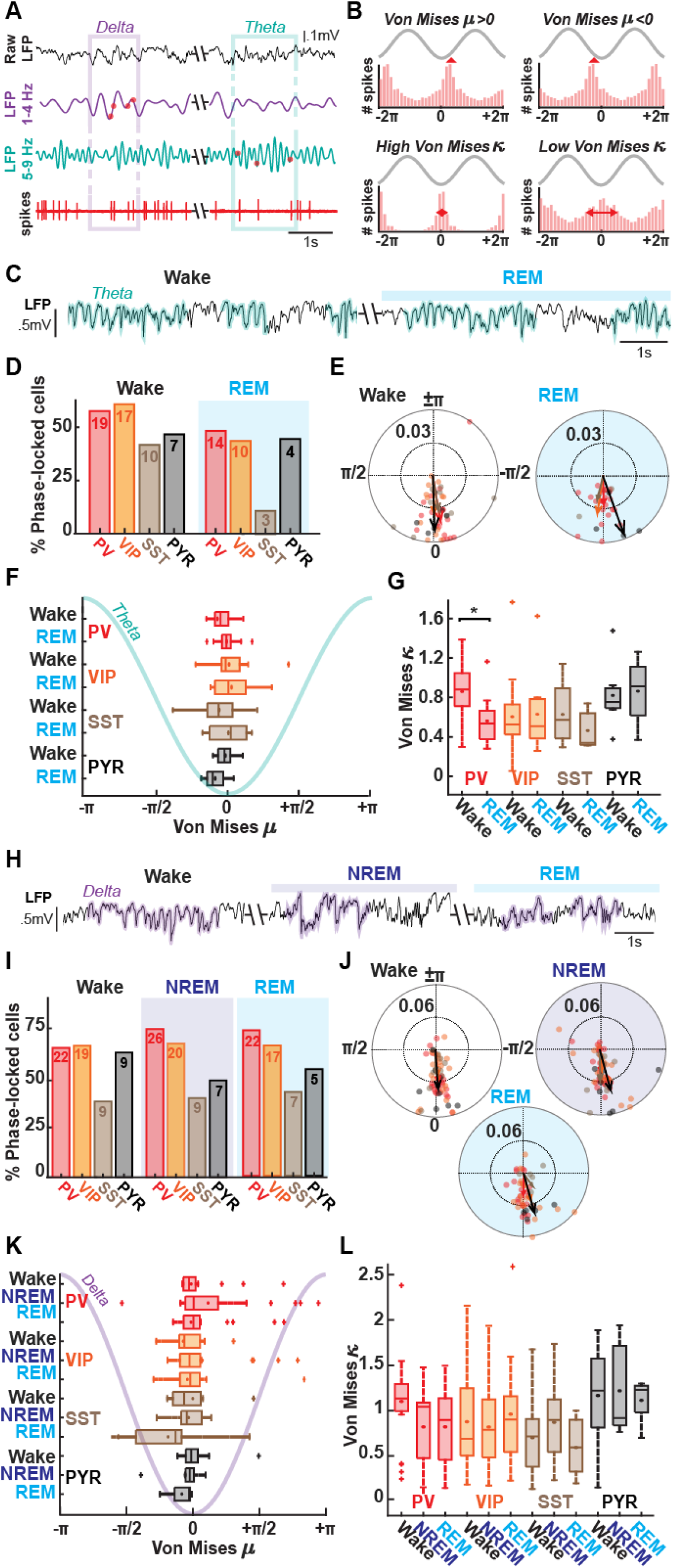
Theta and delta oscillations phase-locking of spikes across vigilance states. **A.** Spike timing on delta (1 - 4 Hz) and theta (5 - 9 Hz) bouts detected on filtered somatosensory barrel cortex (wS1) LFP (see Methods). **B.** Two parameters extracted per cell: the peak μ (on top) and the concentration κ (bottom) of the phase locking computed from a Von Mises curve fitting on spike distribution. **C.** Example raw LFP trace showing detected theta bouts (green) during wake (left) and REM sleep (right). **D.** Percentage of neurons phased-locked to theta oscillations in wake and REM. A Rayleigh test was performed on each cell to detect if they were modulated by theta oscillations (p < 0.01). **E.** Circular plots for all neurons modulated by theta oscillations in wake and in REM. Each dot represents a modulated cell and the arrows the mean for that population (same color code as D). **F.** Average peaks of the phase locking by neuronal subtype. **G**. Average concentration of the phase locking by neuronal subtype. **H.** Example raw LFP trace showing detected delta bouts (violet) during wake (left), NREM (middle) and REM sleep (right). **I.** Same as in D for delta waves detected in wake, NREM and REM sleep. **J.** Same as in E for delta waves. **K.** Same as in F for delta waves. **L.** Same as in G. for delta waves occurring in wake, NREM and REM sleep. *25^th^ and 75^th^ percentiles are represented with median (bar) and mean (dot) in boxplots in F, G, K and L and Kruskal Wallis tests were performed*.

LFP delta oscillations (1-4 Hz) could be observed in all vigilance states, including REM sleep (Funk et al., 2016), although they have been shown to be mostly prominent during wakefulness (Fernandez et al., 2017) and to be interspersed with slow-wave activity during NREM sleep (Crunelli et al., 2006) (Fig. 3*H*). We observed that about 70 % of PV cells were modulated by delta oscillations in wake, NREM and REM (67 %, 76 % and 76 % respectively). A similar proportion of VIP neurons (68 %, 69 % and 68 % for Wake, NREM and REM respectively) but a smaller proportion of SST and PYR cells (SST, 39 %, 40 % and 44 %; PYR, 64 %, and 50 % and 56 % for Wake, NREM and REM sleep respectively) were phase-modulated by ongoing delta activity (Fig. 3*I*).

Similarly to what we observed for oscillations in the theta band, we found no difference amongst modulated cells of any cell type for the position of the peak of the preferred delta phase across all vigilance states (Fig. 3*J, K*). On average, the median concentration median parameter κ was also unchanged for all cell types across all vigilance states (Fig. 3*J, L*).

### Interneurons activity are differentially modulated by spindles

Importantly, we were able to observe clear spindle oscillations during NREM sleep (Fig. 4A). First we investigated whether detected spindles (10 – 17 Hz, see Methods) during NREM sleep modulated the activity of different cortical neurons of S1 barrel layer 2/3. The firing rates of different neuronal subtypes were estimated when spindles occur (Spdl +), and when no spindles were detected on the LFP (Spdl -). On average, firing rates of both PV and PYR cells increased during spindles compared to outside spindles, while the AP activity of SST cells decreased during spindles (PV, Spdl-: 17.1 ± 9.9 Hz; Spdl+: 21.7 ± 12.6 Hz; n = 35, p < 0.001; PYR, Spdl- : 1.5 ± 1.3 Hz; Spdl+: 2.1 ± 2.1 Hz; n = 22, p = 0.010; SST, Spdl-: 3.2 ± 3.2 Hz; Spdl+: 2.8 ± 2.8 Hz; n = 37, p = 0.0017). In contrast, median firing rates of VIP cells remained unchanged by the occurrence of spindle (Fig. 4*B*).

**Figure 4:**
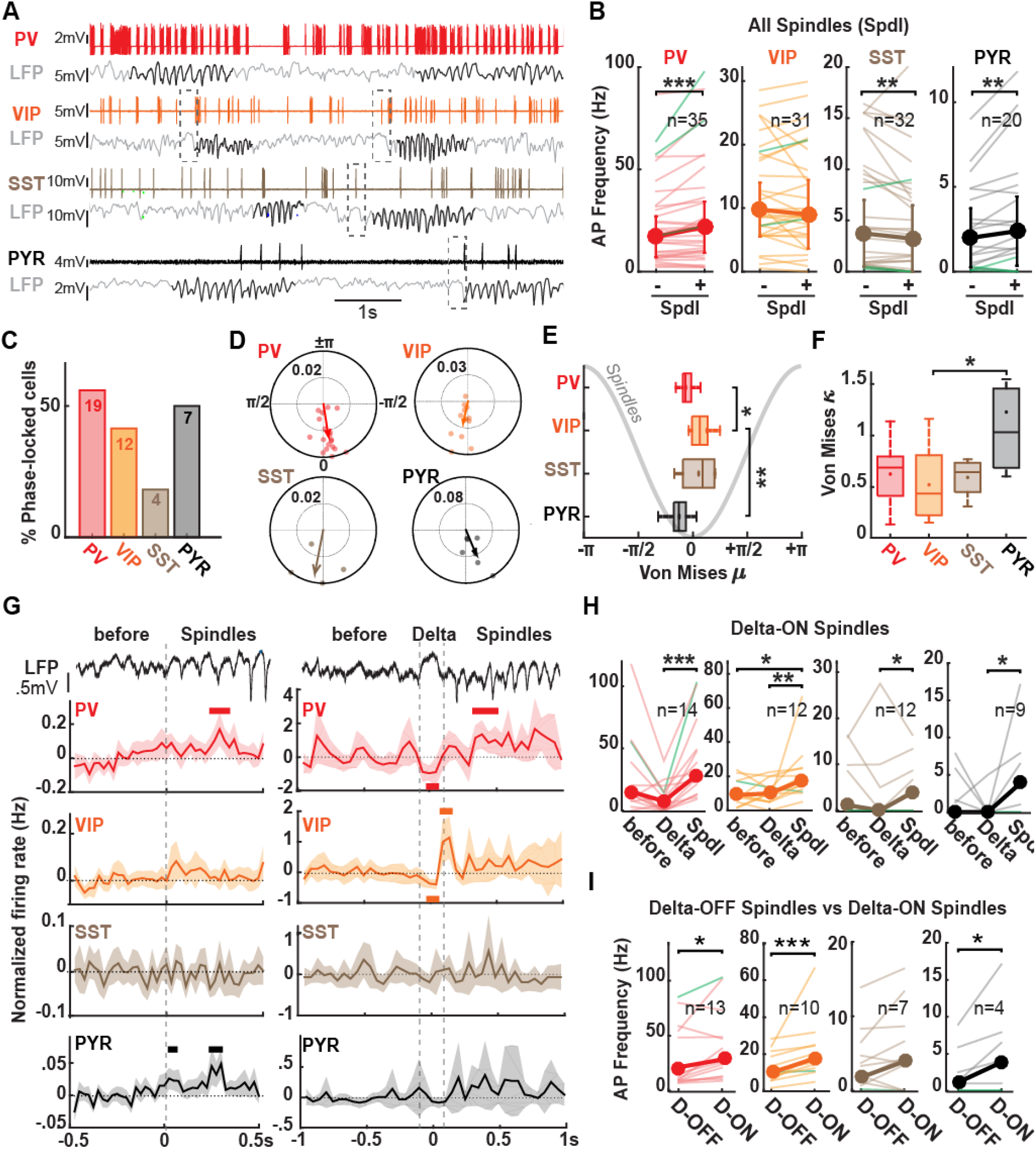
Modulation of interneuronal firing rates by sleep spindles in NREM sleep. **A.** Example raw traces from a PV (top), VIP (middle), SST (middle) and PYR (bottom) neuron together with ongoing LFP activity. Spindle bouts are denoted in black. Delta events occurring before spindle bouts are indicated by dashed grey boxes. **B.** Average AP frequency (bold lines) during spindles (spdl+) and outside spindles (spdl-) for each cell type. Cells recorded in whole-cell patch mode are shown in green. **C.** Percentage of neurons phased-locked to spindle oscillations. A Rayleigh test was performed on each cell to detect if they were modulated by sleep spindles (p < 0.01). **D.** Circular plots for all modulated cells. Each circle represents a modulated cell, and the vector the mean for all modulated neurons. **E.** Average peaks of the phase locking by neuronal subtype. **F.** Average concentrations of the phase locking by neuronal subtype. **G.** Average normalized firing rates across delta-OFF spindles (left) or delta-ON spindles for each cell type. **H.** Average AP frequency before, during Delta event and during spindles for delta-ON spindle bouts for each cell type. **I.** Median firing rates during delta-ON (D-ON) vs delta-OFF (D-OFF) spindle bouts. *Medians and median absolute deviations are represented in B, H and I. Wilcoxon sign rank tests were performed in B and I and Friedman tests were performed in H. 25^th^ and 75^th^ percentiles are represented with median (bar) and mean (dot) in boxplots in E and F and Kruskal Wallis tests were performed. Means and confidence intervals are represented in G*.

We next investigated whether neurons could phase-lock to spindles by performing a Rayleigh test that allowed us to determine whether spiking of individual neurons was significantly locked to a particular phase of a spindle oscillation. A majority (55.9 %) of PV cells were phase-modulated by spindles, as 50.0 % of PYR and 41.4 % of VIP neurons, while only 18.2 % of SST neurons were (Fig. 4*C*). The position of the peak of the preferred phase in radians (rad) was significantly different for VIP cells compared to PV (p = 0.01) and PYR (p = 0.003) (μ, PV: −0.05 ± 0.09 rad; VIP: 0.08 ± 0.14 rad; PYR: −0.11 ± 0.06 rad; *corrected Mann-Whitney test*) (Fig. 4*E*). Indeed, VIP cells discharged 1.2 ms to 2.1 ms after PV neurons and 1.8 ms to 3.0 ms after PYR cells on spindles oscillation (for 10 and 17 Hz spindles respectively). On the other hand, the median concentration parameter κ of PYR cells was higher than the median κ of VIP cells (PYR κ : 1.03 ± 0.4; VIP κ : 0.44 ± 0.26; p = 0.04; *corrected Mann-Whitney test*) (Fig. 4*F*).

In some instances (around 2 %, range: 0 – 18 %), LFP spindles bouts were preceded by a large positive delta wave (1-4 Hz), as previously observed on deeper S1-barrel cortex LFPs during NREM (Urbain et al., 2019) (Fig. 4*A*). These delta waves are thought to correspond to periods of cortical quiescence (Contreras and Steriade, 1995; Maingret et al., 2016). We quantified normalized firing rates before, during delta events when they occurred (Delta), and during spindles (Spdl) and examined how the presence or absence of such delta waves influenced their dynamics. On average, all neuronal subtypes displayed increased firing rates during spindles compared to delta events (PV, Before: 15.0 ± 7.2 Hz; Delta: 7.5 ± 7.5 Hz Spdl: 28.3 ± 16.5 Hz; n = 14, p < 0.001 between Delta and Spdl; VIP, Before: 9.0 ± 5.5 Hz; Delta: 10.0 ± 5.0 Hz; Spdl: 17.6 ± 6.7 Hz; n = 12, p = 0.02 between Before and Spdl, p = 0.004 between Delta and Spdl; SST, Before: 1.1 ± 1.1 Hz; Delta: 0.0 ± 0.0; Spdl: 3.9 ± 3.9; n = 12; PYR, Before: 0.0 ± 0.0 Hz; Delta 0.0 ± 0.0 Hz; Spdl: 3.8 ± 3.8 Hz; n = 9, p = 0.02 between Delta and Spdl) (Fig. 4*H*). Interestingly, the overtime analysis showed us that the increase activity of VIP cells specifically appears at the end of the delta event (Fig. 4*G*).

We then wondered whether the neuronal activity could differ during a spindle, depending on whether or not it was preceded by a delta event. Results showed that all neuronal cell types - except SST cells, displayed a stronger firing rate during spindles preceded by a delta event (PV, Spdl: 13.1 ± 5.0 Hz Delta-Spdl: 28.3 ± 16.5 Hz; n = 14, p = 0.013; VIP, Spdl: 10.8 ± 5.1 Hz; Delta-Spdl: 17.6 ± 6.7 Hz; n = 12, p < 0.001; PYR, Spdl: 0.9 ± 0.9 Hz; Delta-Spdl: 3.8 ± 3.8 Hz; n = 9, p = 0.03) (Fig. 4I).

### Whisking episodes can occur during REM sleep

One of the most striking behavioral features of rodents, which they mainly display during exploration of their environment, is their ability to actively sweep their whiskers back and forth (Welker, 1964). Interestingly, we observed long-lasting (> 1 s) C2 whisker movements during both active wakefulness and throughout the time course of REM sleep episodes (Fig. 5*A*). Those bouts were not equally distributed over the time course of a wake nor REM episode, occurring over longer periods of time at the start of a wake episode and towards the end of a REM episode, respectively (Wake % time spent whisking, 1^st^ part: 9.8 ± 8.3 %; 2^nd^ part: 4.5 ± 8.7%; 3^rd^ part 1.5 ± 5.4 %; n = 92, p = 0,003 for 1^st^ vs 2^nd^ part, p < 0.001 for 1st vs 3^rd^ part; p < 0.001for 2^nd^ vs 3^rd^ part; REM % time spent whisking, 1^st^ part: 0.0 ± 4.6 %; 2^nd^ part: 3.6 ± 8.8 %; 3^rd^ part 6.2 ± 3.6 %; n = 70, p < 0.001 for 1^st^ vs both 2^nd^ and 3^rd^ parts) (Fig. 5*B*). Whisking bouts in REM further differed from their wake counterparts in their frequency of protraction/retraction oscillations, with faster but smaller amplitude series of sweeps superimposed over an already large baseline deflection angle of the C2 whisker occurring during NREM sleep (Peak at around 8 Hz for whisking bouts occurring during wake vs a peak at around 15 Hz for whisking bouts occurring during REM sleep, Fig. 5*A, C*), and their median maximal amplitudes (Wake: 46.2 ± 8.1 °; n = 92; REM: 32.8 ± 5.5 °; n = 70, p < 0.001) (Fig. 5*D*). However, their durations were similar to those occurring during wake (Whisking bouts duration, Wake: 1.85 ± 0.97 s; n = 92; REM: 1.76 ± 1.76 s, n = 70) (Fig. 5*D*). Mice therefore display whisking bouts with specific characteristics during REM sleep, but their durations preclude them from being categorized as twitches (Tiriac et al., 2012).

**Figure 5:**
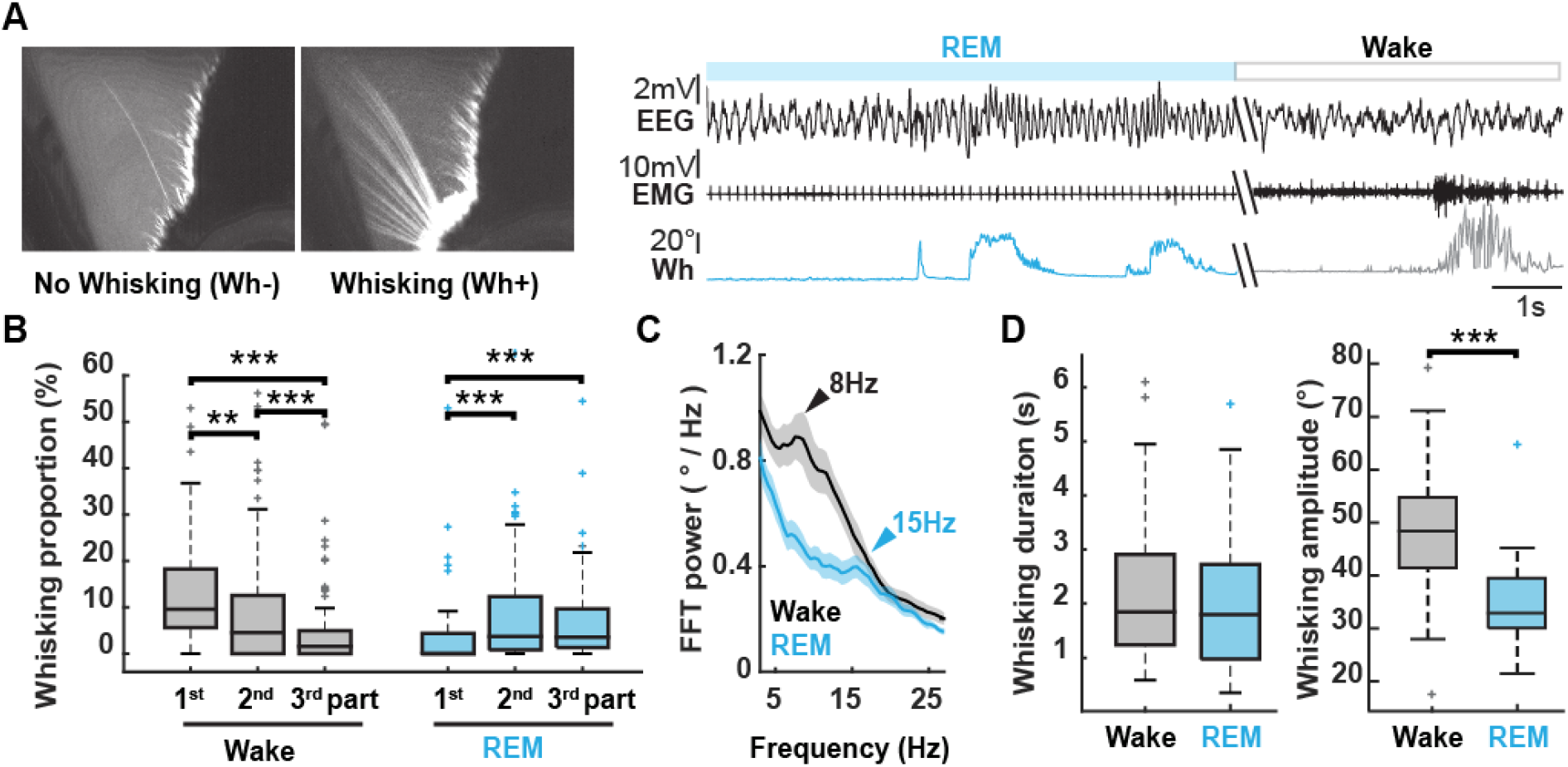
Whisking bouts occur during both wake and REM sleep in naturally-sleeping mice. **A.** Left: Mean picture of C2 whisker observed through a high-speed infrared camera in the absence or presence of whisker protractions. Right, Example EEG (top), EMG (middle), and Whisker position (bottom) traces during REM sleep and wake, showing bouts of whisking (> 1s) in both vigilance states. **B.** Percentage of time spent whisking during the first, second and third part of wake and REM sleep across all recorded episodes. Dots denote results per recording session. **C.** Grand mean average Fast Fourier Transform (FTT) of whisker position during whisking bouts occurring in wake (black) and REM sleep (cyan). FFTs were computed from 1s time windows for all bouts across all recording sessions. A peak at around 8 Hz during wake-whisking and at 15 Hz during REM-whisking are marked by black and blue arrows respectively. **D.** Average whisking bout durations (left) and amplitudes (right) during wake and REM sleep. *25^th^ and 75^th^ percentiles are represented with median (bar) in boxplots in B and D. Friedman tests were performed in B and Wilcoxon sign rank tests were performed in D. Means and confidence intervals are represented in C*.

### Whisking induces changes in interneuronal firing activity during both wake and REM sleep

Previous studies uncovered differential modulations of GABAergic cells neuronal firing during free whisking (Gentet et al., 2010, 2012; Muñoz et al., 2017) or active tactile behavior (Yu et al., 2019). Since we observed whisking behavior in our head-restrained, naturally-sleeping mice, during both wake and REM sleep, we next examined whether it could differentially affect neuronal firing rates in our population of layer 2/3 cortical neurons during these two fundamentally different cognitive brain states (Fig. 6). We quantified the average firing rates of different cellular subtypes when the mouse was whisking (Wh+) versus when no whisking occurred (Wh-). Furthermore, twitches were excluded from the analysis (seeMethods). We observed different pattern of activity across different cells in both loose-patch (Fig. 6*A*) and whole-cell patch-clamp (Fig. 6*B*) configurations. On population data, AP firing in both PV and PYR neurons was not modulated by whisking bouts occurring during either wake nor REM sleep. On the other hand, VIP cells firing activity was increased during whisking in both wake and REM sleep (Wake Wh-: 9.4 ± 6.1 Hz; Wake Wh+: 19.0 ± 9.9 Hz; n = 28; REM Wh-: 10.7 ± 5.4 Hz; REM Wh+: 13.4 ± 6.3 Hz; n = 23, p < 0.001 for whisking effects in Wake, p = 0.01 for whisking effects in REM). In contrast,SST cells significantly decreased their AP firing whisking in wake (Wake Wh-: 4.0 ± 3.2 Hz; Wake Wh+: 3.2 ± 3.0 Hz; n = 27 p = 0.02) and alsotend to decrease during whisking bouts in REM sleep (REM Wh-: 6.3 ± 5.4 Hz; REM Wh+: 4.9 ± 4.9 Hz; n = 15) (Fig. 6*C*). These results on population averages may not however reflect the behavior of individual neurons during whisking throughout the sleep-wake cycle. Therefore, in order to better understand whether whisking could differentially affect the firing rates of different neurons, we subtracted individual neuronal firing rates during whisking by the firing rates during non-whisking, divided by the firing rate during non-whisking, to obtain an index as a % change for each cell. We defined cells as a whisker-active (Wh+) or whisker-inhibited (Wh-) when the absolute percentage of change exceeded +15 % and −15 %, respectively. Half of PV cells were inhibited during whisking in wakefulness (53 %) while 29% were activated. During REM sleep however, PV neurons activity remained for half of them unchanged (53 %). In contrast, in both wake and REM sleep, a big majority of VIP cells displayed an increased firing rate when whisking was observed (83 % and 67 % respectively) and only a small part of VIP neurons were inhibited (11 % and 22 % respectively). (Fig; 6*D*). In other hand, most of SST neurons are inhibited by whisking in wake and REM sleep (56 % and 44 % respectively), such as PYR cells (67 % and 78 % respectively). (Fig; 6*D*). Finally, we compared the percentages change in wake and REM for each individual neuron grouped by subtype. Thanks to this analysis, we were able to known if one specific neuron adopted a different firing rate modulation in wake compared to REM sleep. In our population of VIP cells, this percentage change dropped significantly in REM compared to wake, indicating that the increase in VIP cells AP firing that occurs during whisking in wake is larger than the increase observed during REM sleep (VIP Wake % change: +78.5 ± 69.4 % vs +22.7 ± 23.6 % in REM; n = 18, p = 0.01). For PV, PYR and SST cells, the percentage changes remained stable on average in wake compared to REM (Fig. 6*E*). In summary, VIP cells population is mostly homogeneous, with a strong increased activity during whisking in wake and a smaller one during REM sleep, while PV, SST and PYR neurons display a more heterogeneous pattern.

**Figure 6:**
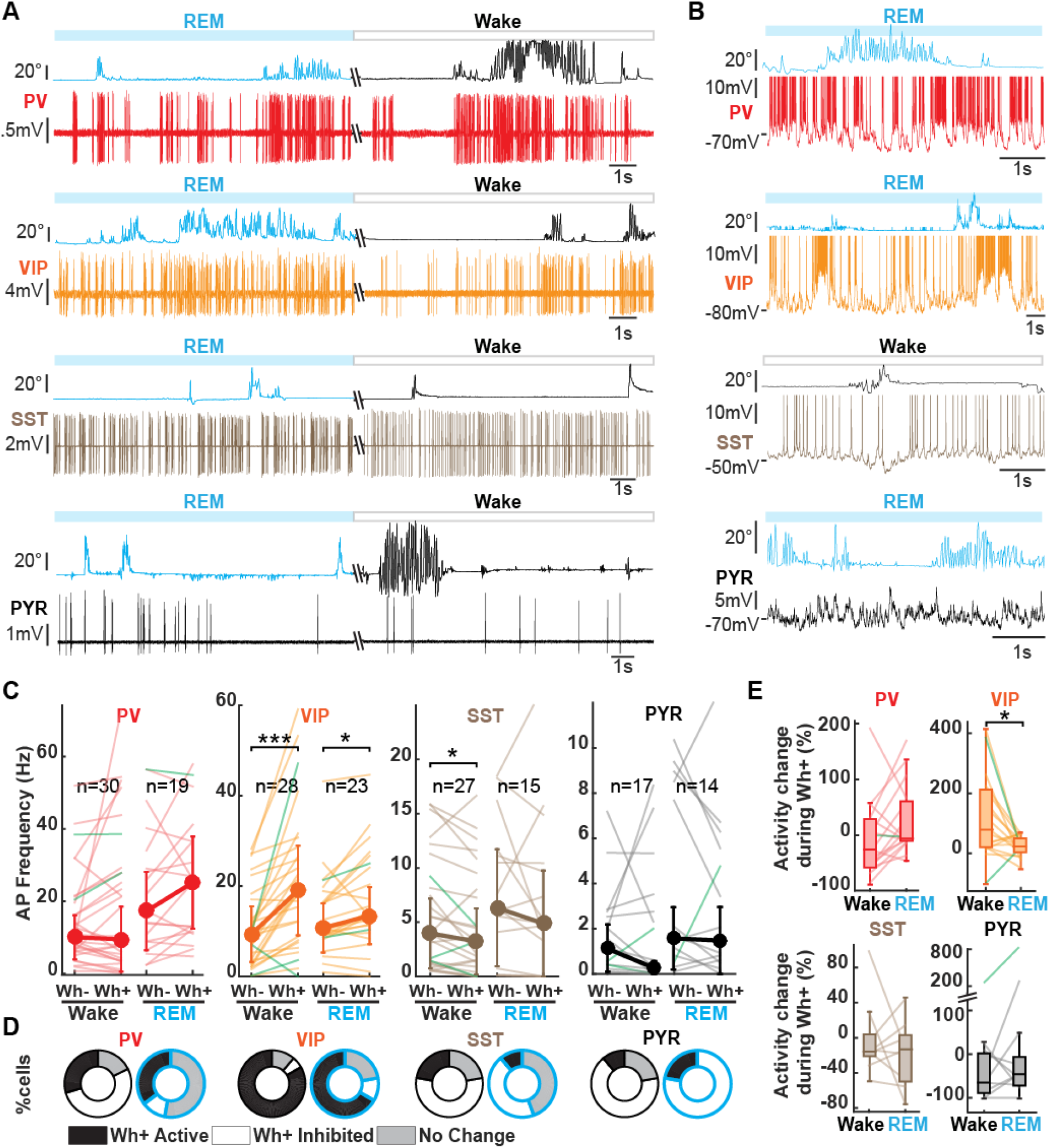
Whisking modulates cortical interneurons firing rates. **A.** Example raw traces of loose-patch recordings and whisker movements in wake and REM sleep for one PV (top, red), one VIP (middle, orange), one SST (middle, beige) and one PYR (bottom, black). **B.** Example raw traces of whole-cell recordings and whisker movement in wake or REM sleep (same color code as A). **C.** Median firing rates for PV, VIP, SST and PYR neurons during whisking bouts (Wh+) vs no whisking (Wh-). Cells recorded in loose-patch are represented in light color and whole-cell patch-clamped neurons in green. **D.** Percentage of cells per subtype and per vigilance state displaying either increased, decreased or no change in their firing rates during whisking vs no whisking. **E.** Activity change in % during whisking in wake vs REM compared to no whisking in wake and REM sleep. *Medians and median absolute deviations are represented in C. 25^th^ and 75^th^ percentiles are represented with median (bar) in boxplots in E. Wilcoxon sign rank test were performed in C and E*.

## DISCUSSION

Cortical activity throughout the sleep/wake cycle needs to be highly dynamic, in order to perform different behavioral tasks during wakefulness and allow the consolidation or removal of unnecessary memory traces during sleep (Kay and Frank, 2019). Local GABAergic interneurons can efficiently control the spike timing of principal neurons via their precise targeting of different regions of the somato-dendritic arborizations of pyramidal cells (Gentet, 2012). Layer 2/3 of the somatosensory barrel cortex (wS1) is involved in the processing of whisker sensory information and can perform computations of both simple and complex tasks in a goal-oriented manner (Petersen, 2019). While PV cells primarily exert perisomatic inhibition onto excitatory cells (Kawaguchi and Kubota, 1997), SST cells, comprised essentially of Martinotti-type neurons (Jiang et al., 2015) control the dendritic encoding of synaptic inputs in the somatosensory cortex (Murayama et al., 2009; Gentet et al., 2012). How local GABAergic inhibitory cells regulate this circuit during different vigilance states of the sleep/wake cycle, and how ongoing brain oscillations or whisking behaviors modulate their firing patterns or activity, remains however poorly understood.

In this study, we first performed calcium-imaging of neuronal population activity in layer 2/3 of wS1 in different transgenic mouse lines where genetically-identified subpopulations of GABAergic neurons could be visualized. This initial study first revealed that PV cell activity is increased during both NREM and REM sleep stages, VIP activity is increased specifically during REM sleep, and SST and PYR neuronal activity remains overall stable over the time course of the sleep/wake cycle. These findings share both similarities and differences with a previous study in the motor cortex, showing elevated PV cell activity during REM, but reduced activity of PV, SST and PYR neurons during NREM compared to wakefulness (Niethard et al., 2016). Such discrepancies could be explained by differences in the brain region under investigation, or experimental procedures. In our study, mice were habituated to the head-fixation device for considerably longer periods of time than in the study by Niethard and Colleagues (3 weeks in the present study versus one day before imaging sessions began). Our animals might therefore have displayed reduced levels of stress from the head-restraint during imaging sessions. In any case, our results under calcium-imaging were confirmed by our targeted recordings of individual neurons in loose-patch or whole-cell patch-clamp configurations. Such recordings also offer the advantage of a more precise temporal resolution of AP firing than calcium-imaging, and allowed us to simultaneously record the LFP by placing a glass pipette within layer 2/3 of the same column from which our targeted single-unit recordings originated (Perrenoud et al., 2016). Using this combination of recording elements, we first confirmed our findings that overall activity of PV cells is significantly increased in both NREM and REM sleep compared to wake, and that VIP cells are specifically more active during REM sleep. These measures of cell type-specific average firing activity according to vigilance states offer a good indicator that the excitatory/inhibitory balance is highly dynamic across the sleep/wake cycle. In our study, we further investigated those dynamics along transitions between vigilance states and we found that PV activity rapidly decreased at wake onset from NREM and REM sleep episodes, as previously reported for transitions from NREM to wake for fast-spiking, putative PV neurons in the mouse frontal cortex (Miyawaki et al., 2019). This rapid decrease could be due to the rapid activation of the ascending arousal system. For example, wake-specific histaminergic neurons of the tuberomamillary nucleus and norepinephrine neurons of the locus cerruleus, both with extensive projections throughout the cerebral cortex, have been shown to rapidly activate at wake onset (Takahashi et al., 2006, 2010).

Notably, we found that distribution of pyramidal cell activity broadened during REM sleep as described by Miyawaki et al. (2019). We therefore decided to split pyramidal cell population into two subgroups on the basis of their basal activity during REM. While activity at the population level remained stable across vigilance states, this separation uncovered subtler changes in the activity of principal neurons throughout the sleep/wake cycle. The neurons with the highest basal levels of REM activity were significantly less active during wake and NREM sleep, while neurons with low REM activity did not display any significant overall changes in AP firing frequency across vigilance states.

Interestingly, while the average AP activity of SST neurons remained stable across the sleep/wake cycle, their firing patterns, as well as those of other types of interneurons, shifted to a more burst-like behavior during NREM sleep compared to wake. This shift could be explained by the more synchronized state of cortex during NREM sleep, with the time windows for neuronal firing being temporally-restricted for all neurons compared to wake. However, the firing irregularity of GABAergic neurons was differentially modulated during NREM sleep, with increased firing irregularity being present in VIP neurons but not PV nor SST cells. Thus, VIP neurons may play a major in the re-organization of GABAergic neuronal activity patterns during NREM, through its strong disinhibitory influence over local cortical interneurons (Pfeffer et al., 2013). The important role of VIP neurons firing dynamics during NREM was further evidenced by their firing behavior during sequences of delta waves-spindles. Delta waves preceding spindles have recently been shown to play a role in memory consolidation via the recruitment of local cell assemblies performing isolated cortical computations (Todorova and Zugaro, 2019). Interestingly, we observed subtle temporal changes in the activity of GABAergic VIP neurons over the time course of LFP delta – spindles sequences that suggest that the disinhibitory control exerted by this interneuronal subpopulation is reduced during delta events, but rises quickly towards the end, and at the time of emergence of spindle oscillations (Fig. 6 *H*). This could lead to short, but temporally precise time windows, during which removal of inhibition onto excitatory cell assemblies would allow cortical replay sequences to be carried out (Ji and Wilson, 2007).

During NREM sleep, we further found an increased activity during periods of LFP spindles in both PV and excitatory layer 2/3 pyramidal neurons, but a decrease in SST overall neuronal activity, as previously reported across different cortical regions (Niethard et al., 2018). In addition, we report than amongst neurons that were modulated by spindle oscillations, PV and pyramidal cells fired at a preferentially earlier phase than their VIP counterparts, indicating that these cells might contribute to the emergence and/or amplification of these LFP oscillations through their sequential activation. While spindles are considered to be thalamic in origin, and induce phase-locking of rhythms in both neocortical and hippocampal networks (Latchoumane et al., 2017), it is currently unclear whether the spindles that we recorded on our LFP electrodes are mainly a representation of these propagating spindles, an emanation of local circuit properties, or a combination of both, as LFPs are notoriously difficult to interpret (Einevoll et al., 2013).

A majority of local cortical neurons were modulated by ongoing LFP slow delta waves during both wake and sleep. Only SSTs neurons displayed relatively weaker modulation, in accordance with a previous finding suggesting that their membrane potential dynamics were not correlated to those of surrounding neurons during quiet wakefulness (Gentet et al., 2012). Interestingly, this weaker modulation remained stable throughout the sleep/wake cycle as well as the preferred phase locking of all cortical neurons.

Whisking behavior in rodents is associated with their nocturnal foraging habits, as they explore their environment through the whisker sensing of surrounding objects (Petersen, 2014). While during REM sleep, mice display, just like humans, rapid-eye movements (Fulda et al., 2011), we surprisingly observed long-lasting bouts of whisking behavior in mice during REM sleep, despite the complete muscle atonia. We distinguish these bouts of whisking from previously reported REM “twitches” in the rat neonate (Tiriac et al., 2012) by their durations (over 1 second versus a few tens of milliseconds for twitches) and the amplitude of their protraction. Nevertheless, they differed from their wake counterparts in their higher peak frequency of oscillations, corresponding to smaller but faster protraction/retraction sweeps from an already protracted whisker position from rest. In this paper, we first confirm results obtained from previous studies showing whisking-related changes in interneuronal AP firing in awake head-fixed mice: while PV cells displayed heterogeneous modulation of their activity following the onset of free whisking, VIP neurons were more consistently activated during whisking, while SST firing was reduced, as previous findings (Gentet et al., 2010, 2012; Lee et al., 2013; Muñoz et al., 2017). During REM sleep however, whisking induced smaller increases and no significant decrease in VIP and SST neuronal activity, respectively. This differential modulation of VIP neurons and their target SST cells could help isolate the local circuit from top-down cortico-cortical and thalamic afferents impinging onto distal dendrites of excitatory principal pyramidal cells (Oda et al., 2004; Larkum et al., 2009). This raises the interesting prospect that REM whisking bouts might correspond to a brain state during which internalization of awake whisking behavior is re-experienced.

In summary, our study sheds light on the complex interplay between local cortical interneuronal firing activity and ongoing brain oscillatory dynamics across different vigilance states. Especially, we have shown that PV and VIP neurons may play prominent roles in dynamically suppressing, disengaging, or activating the local circuits at important time points of the sleep/wake cycle. The functional consequences of this dynamic interplay might be better observed at the level of perception of sensory inputs during sleep (Velluti, 1997; Nir et al., 2015) or consolidation of sensory memory traces acquired during wake (Buzsáki, 1989; van Dongen et al., 2011).

## Author contribution

L.G. designed research; A.B performed research; A.B analyzed data; M.B. and L.G helped in data analysis; A.B. and L.G. wrote the manuscript; N.U. commented on the manuscript.

## Acknowledgements

We would like to thank J.C. Comte for technical assistance. This work was funded by a doctoral studentship from the French Ministry of Education (A.B.), ANR PARADOX (ANR-17-CE16-0024) (A.B. and L.G.), and FLAG-ERA JTC 2015 project CANON (co-financed by ANR) (M.B. and L.G.).

